# Classification of bacterial nanowire proteins using Machine Learning and Feature Engineering model

**DOI:** 10.1101/2023.05.03.539336

**Authors:** Dheeraj Raya, Vincent Peta, Alain Bomgni, Tuyen Du Do, Jawaharraj Kalimuthu, David R. Salem, Venkataramana Gadhamshetty, Etienne Z. Gnimpieba, Saurabh Sudha Dhiman

## Abstract

Nanowires (NW) have been extensively studied for *Shewanella* spp. and *Geobacter* spp. and are mostly produced by Type IV pili or multiheme c-type cytochrome. Electron transfer via NW is the most studied mechanism in microbially induced corrosion, with recent interest in application in bioelectronics and biosensor. In this study, a machine learning (ML) based tool was developed to classify NW proteins. A manually curated 999 protein collection was developed as an NW protein dataset. Gene ontology analysis of the dataset revealed microbial NW is part of membranal proteins with metal ion binding motifs and plays a central role in electron transfer activity. Random Forest (RF), support vector machine (SVM), and extreme gradient boost (XGBoost) models were implemented in the prediction model and were observed to identify target proteins based on functional, structural, and physicochemical properties with 89.33%, 95.6%, and 99.99% accuracy. Dipetide amino acid composition, transition, and distribution protein features of NW are key important features aiding in the model’s high performance.

## 1. Introduction

Extracellular electron transfer (EET) between microbial communities and metal through direct electron transfer via protein (nanowires) or indirect electron transfer mediated by electron shuttles is the crucial fundamental mechanism that has been studied for microbially induced corrosion (1). Microbial nanowires are electrically conductive bacterial extensions allowing electron transport over long distances. Nanowires have useful impacts in bioenergy, bioremediation, and bioelectronics and play a significant part in the carbon and mineral cycle in anaerobic soils and sediments and novel methods for interspecies energy exchange (2).

Many microbes have been reported to be involved in EET and nanowires’ formation; however, *Shewanella oneidensis* (3,4) and *Geobacter sulfurreducens* (5) have been studied in depth. Most protein nanowires are electrically conductive pili assembled from PilA, type IV pili, and multiheme c-type cytochrome such as OmcS and OmcZ in *G. sulfurreducens*. Only a few studies have reported nanowires in sulfate-reducing bacteria (SRB) (6,7) that contribute towards microbially induced corrosion in aqueous systems.

The general empirical criteria that have been successful in predicting whether pilin genes may yield proteins that assemble into e-pili are (1) aromatic amino acids located in the same position along the pilin sequence as aromatic amino acids thought to be required for conductivity in *G. sulfurreducens* e-pili; (2) aromatic amino acid abundance of 9% or greater in the mature pilin sequence; and (3) no regions along the pilin of more than 40 amino acids without aromatic amino acids (7). However, generalized empirical criteria of microbial nanowires still need to be included.

Using existing data, machine learning (ML) has established itself as a powerful, efficient tool to make predictions on various contexts, including antibiofilm properties, quorum sensing molecules, and many more. An experimentally validated peptide sequence was used to construct an ML model to predict antibiofilm peptides with an accuracy of 97.83% (8). Similarly, Srivastava et al. (2020) used 122 molecules known for antimicrobial properties from either plant, human, or synthetic origin as a positive dataset. They included 189 metabolites as a negative dataset to train an ML model such as random forest and support vector machine to successfully predict a set of independent data for biofilm inhibitory properties (9). Hence, protein-based sequence extraction and an ML approach would help predict the properties of proteins. Sequence-based protein classifiers assign labels based on a set of features and real numbers, capturing some specific sequence property. However, features extraction and appropriate features would enable researchers to predict better the protein’s functionality in question (10). Several protein features, such as amino acid composition, dipeptide composition, property-based profile features, and amino acid distance-based features, are used to classify the protein based on these sequence features.

This study created a manually curated dataset of nanowire proteins from available databases (Uniprot and NCBI). The database consisted of proteins from *G. sulfurreducens* and *S. oneidensis* capable of electron transfer when assayed *invivo* or in-vitro. This study aims to develop a prediction model for the classification of nanowire proteins and assess the performance of different algorithms. Secondly, gene ontology analysis was performed to determine microbial nanowire proteins’ cellular localization and molecular function from the database. Finally, feature engineering was implemented to assess the feature subset’s contribution to the model’s performance and their contribution.

## 2. Material and Methods

### 2.1. Preparation of dataset

Proteins involved in nanowire biogenesis were searched through several publicly available databases to extract associated sequences. Initially, the Uniprot database was searched using the keyword ‘nanowire,’ resulting in many redundant entries (11). Eight hundred forty-two proteins were extracted following manual curation to ensure the quality of extracted protein entries. Similarly, following manual curation of access from the NCBI (https://www.ncbi.nlm.nih.gov/) database using the same keyword, 157 proteins were extracted for further analysis. A total of 999 proteins were retrieved to prepare the positive dataset. There is no general rule of thumb for the selection of negative datasets, and usually, negative datasets are selected such that these datasets are not overlapping with the positive dataset. The dataset was randomly divided into training and test datasets using a ratio of 33:67.

#### Protein features extraction and composition

Twenty-one types of feature vectors were extracted using iFeature (12). Amino acid composition descriptors groups, mainly amino acid composition (AAC), dipeptide composition (DPC), and tripeptide composition (TPC) features, were extracted to represent the percentage of each amino acid in protein sequences. Amino acid composition and dipeptide composition generates 20 and 400 dimensions. This dimension specifies the number of occurrences with either the total number of residues in protein or dipeptides in protein.

Composition, transition, and distribution descriptors group (mainly CTDC, CTDD, CTDT) were extracted for numerical representation of peptide physicochemical properties. Physicochemical properties of amino acids describe the parameters such as oxygen, sulfur, nitrogen, hydrogen, and carbon content and describe amino acid acidity and basicity, size, polarity, and charged percentage (13). A detailed list of protein descriptors used in this study is provided in Table S1. In total, 19,856 features were extracted for a sequence of proteins.

### 2.2. Features preprocessing and Machine Learning model

ML model quality and precision depend on the data quality used for training. Redundant features, such as constant features having the same values and quasi-constant features, were removed from the extracted features. After processing, features were used to train the ML model. The prediction model was built using several ML algorithms, including support vector machine (SVM), Random Forest (RF), and Extreme gradient boosting (XGBoost). SVM is one of the most commonly used classifiers in peptide prediction, which works particularly well for binary classification problems. SVM is a robust model that can be used for classification and regression (14).

RF is an ensemble of decision trees, where every tree learns the classification on a random subset of the domain pairs. The most discriminating descriptor for each tree is picked from a randomly chosen subset of m descriptors, where m is substantially less than the total number of descriptors at each splitting or decision node (15). XGBoost is used for supervised learning and has excellent scalability and a high running speed (16). A five-fold cross-validation of the training dataset was performed to address the potential overfitting problems. The whole dataset was divided into five sets such that in each round, four sets were used in training, and one was set aside for testing. Once the classification models were ready, their performance was tested in terms of accuracy and precision.

### 2.3. Gene ontology enrichment

The positive dataset was categorized based on gene ontology terms (biological process, molecular function, and cellular component) using the InterPro database and UniProt (11,17). R statistical language was used for the computational analysis and generation of figures.

## 3. Results and Discussion

### 3.1. Analysis and characteristics of proteins in the positive dataset

The positive (nanowire proteins) and negative datasets were meticulously curated for machine learning and subsequent model development. Positive datasets were manually curated to include either type IV pili, Nanowire_3heme (IPR026352), and multihaem_cyt_sf (IPR036280) family domain. IPR026352 domains bind three hemes and occur as repeating units in GSU_1996, whose crystal structure shows elongation and which functions in extracellular electron transport processes. Similarly, IPR036280 domains in GSU_2731 contain multiple CxxCH motifs involved in the extracellular electron transfer process. A total of 999 positive datasets were manually curated from NCBI and Uniprot for this study. OmcS, OmcZ, mtrA, and PilA were reported to be active in extracellular electron transfer and nanowire development in *G. sulfurreducens* and *S. oneidensis was* part of the positive dataset.

The positive dataset used in this study comprises a large number of sequences with similar lengths ranging from 50 to 350 amino acid residues (Fig. 1A). Type IV pili, such as PilA, which are involved in nanowire development and electron transfer, range from 60 - 90 amino acids (18). In contrast, multiheme-based nanowires, such as OmcS, OmcC, and OmcZ, have lengths of more than 200 amino acids (19). While analyzing the distribution of 20 amino acids in proteins of the positive dataset, significantly higher counts of alanine (A), lysine (K), glycine (G) and lower counts of tyrosine (Y), glutamine (Q), and tryptophan (W) were observed (Fig 1B). These results suggest that the positive dataset is enriched in the positively charged amino acid residue, consistent with the previous report that nanowires originating from type IV pili and OmcS have positively charged pockets that bind to negatively charged Fe(III) oxides during reduction (20,21). Upon analyzing the physicochemical properties of the positive dataset, it was observed that a significantly lower fraction of polar residues with hydrophobic amino acid residues were present (Fig 1C). This observation is consistent with a previous report indicating that cytochrome-based nanowires are predominately hydrophobic (22).

**Figure 1.**
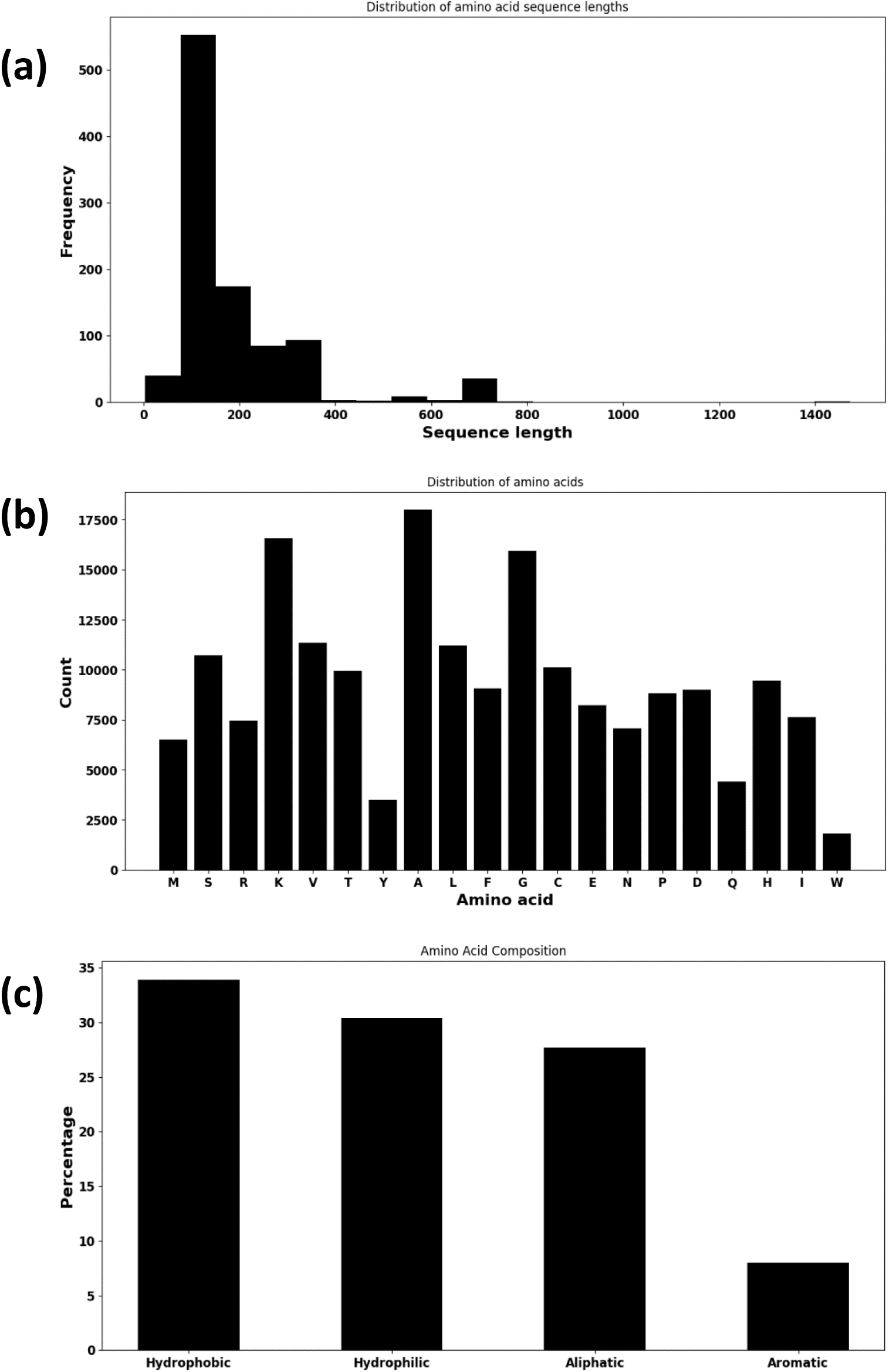
Sequence properties of NW forming protein (a) extracted from UniProt and NCBI database, and (b,c) amino acid composition

### 3.2. Electron transfer activity and metal ion binding in nanowire

For gene ontology (GO) classification of the positive dataset, biological process (BP), cellular component (CC), and molecular function (MF) were used. BP represents protein-based pathways, MF for molecular functions, and CC for the position of a protein in relation to a cellular component or structure. GO terms associated with each protein in the positive dataset were unavailable through Uniprot. Hence, GO terms associated with 3.43% of the positive dataset were retrieved and analyzed.

Electron transport chain was the primary term related to BP. However, only four proteins in the positive dataset had associated BP terms and were not used in further analyses. CC enrichment analysis revealed the involvement of membranal proteins (Fig 2). Membrane proteins like outer membrane cytochrome proteins and type IV pili are the first or final step in the electron transfer via nanowires. For instance, nanowires produced by *S. oneidensis* or *G. sulfurreducens* are localized in the outer membrane (21,23). Localization of PilA or cytochrome proteins in the outer membrane is the initial step in the biogenesis of microbial nanowires and can greatly affect the functionality of a nanowire (4).

**Figure 2.**
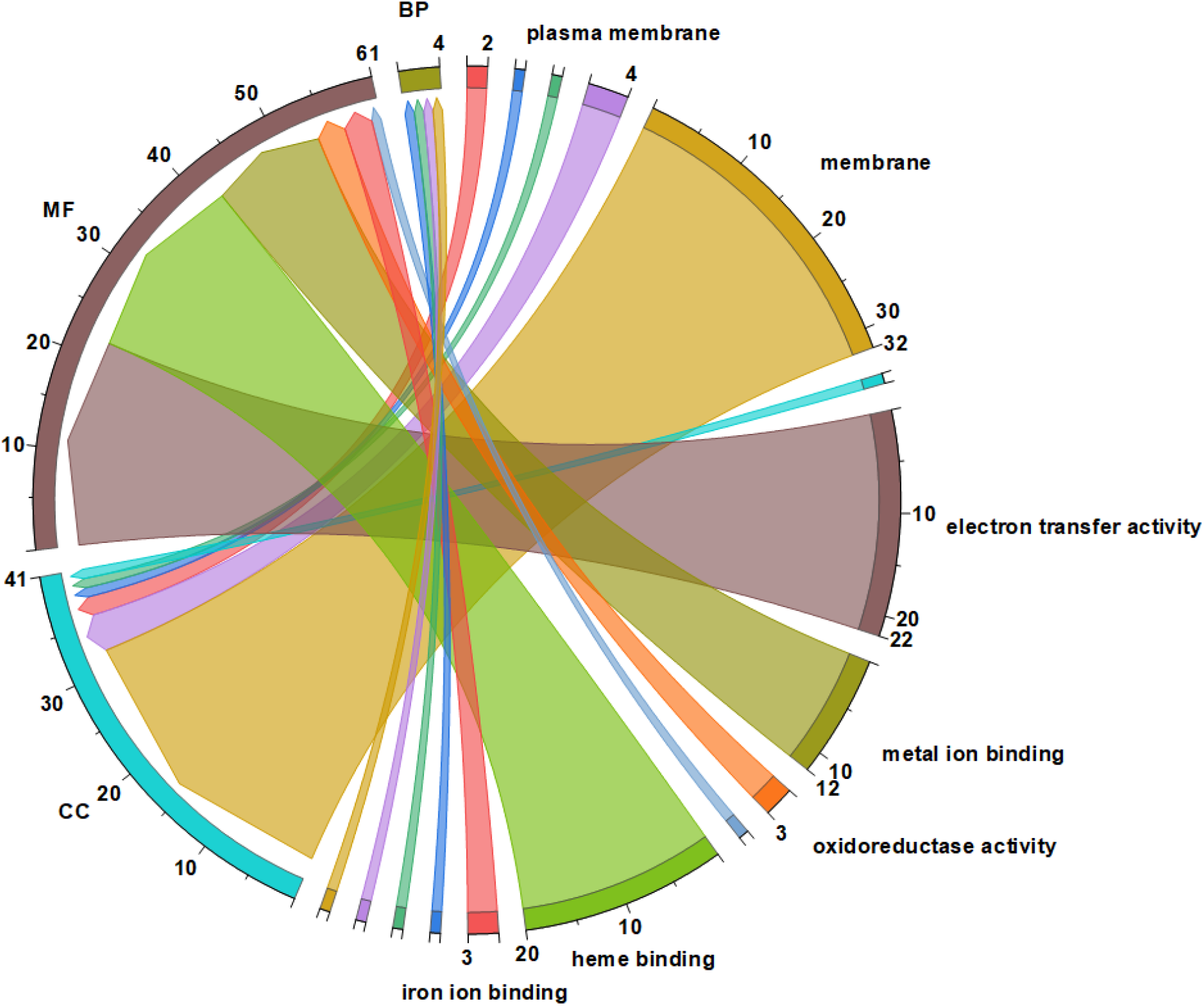
Gene ontology (GO) of the associated nanowire protein dataset. *BP: Biological process; CC: Cellular Component; MF: Molecular Function*

With regards to MF analysis, the three most enriched terms were electron transfer activity (GO:0009055), metal ion binding (GO: 0046872), and heme binding (GO: 0020037). Electron transfer activity is the key feature of a microbial nanowire protein. Cytochromes such as OmcS, OmcZ, or type IV pili mediate electrons between the bacterial cell and extracellular electron acceptors or insoluble metal oxides. Metal ion binding is another important feature of microbial NW, aiding in binding with insoluble metal oxides. Both OmcS and PilA proteins contain positively charged domains that can bind with negatively charged metal oxides (21,24). The interaction between metal oxides and nanowire proteins helps to facilitate the extracellular electron transfer activity when an insoluble electron acceptor is present. Finally, OmcS and OmcZ are involved in nanowire formation and contain multiple heme groups. For instance, each OmcS monomer accounts for 6 heme-binding motifs, whereas OmcZ has eight heme-binding motifs (25,26). The arrangement of heme groups in a linear fashion, with each heme group linked to the next in series, is the mechanism of electron transfer in a multiheme-based nanowires (27). As nanowire formation occurs during the limitation of electrons, at least two major roles of nanowire proteins should involve electron transfer activity and metal ion binding.

### 3.3. Machine learning model based on subset of features

Based on the twenty-one features extracted, RF models were generated using each feature as input. The models achieved an accuracy greater than 94.21% for each subset of features, with the Ctriad feature giving maximum accuracy (Fig 3). The moran, SOCNumber, Geary, and Auto covariance (AC) showed a comparatively lower accuracy of approx. 94%. Similar trends were observed for each input feature regarding precision (Table S1). Amino acid composition descriptors group (AAC, CKSAAP, DPC, and TPC), Composition, transition, distribution descriptor group (CTDC, CTDT, and CTDD), Amino acid Index protein descriptors, Z-scale protein descriptors showed higher accuracy ranging in 98-99%. Amino acid composition-based descriptors have been previously used to classify the subcellular localization of proteins (28). Composition, transition, distribution descriptors group, and amino acid index protein descriptors have been widely used to predict the protein functions (29).

**Figure 3.**
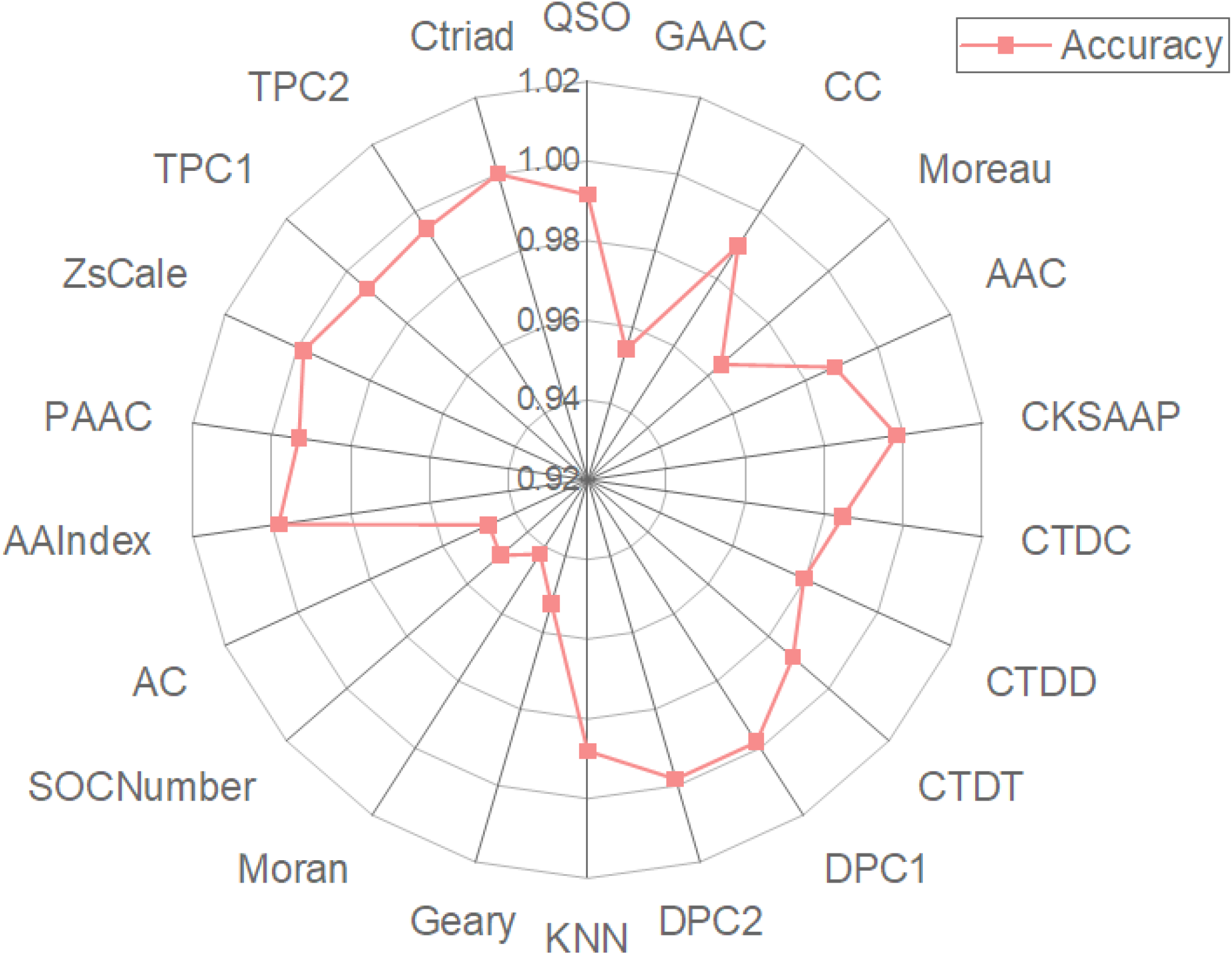
Accuracy of random forest model using each input descriptor as features.

### 3.4. Model evaluation with different classifiers

The primary and secondary structure information of proteins and their functional properties were used to develop ML models to identify and understand the features unique to nanowire proteins. 19,859 features obtained from twenty-one protein descriptors for a single sequence were used to train the ML models. The radial basis kernel was used without recursive feature elimination in the SVM model. The radial bias kernel in SVM performed well and achieved an accuracy of 89.31% (Fig S1). For XGBoost and RF models, the model parameters, number of estimators, and maximum depth were tuned. The performance of the models is presented in Fig S1. The accuracy of the RF model was 95.61%, while that of XGBoost was similar.

In contrast to the single input feature-based ML model, the performance ML model decreased using a combination of all features. In reality, training the machine learning model utilizing high dimensional features tends to behave poorly, and this phenomenon is called the “curse of dimensionality” (30). Important features were considered input features to remove redundant features and improve the model’s performance. Hence, using the top hundred important features to reduce the dimensionality of a model, RF model accuracy increased to 95.77% (Fig 4). The RF model produced a good prediction performance, showing that the features can effectively characterize nanowire proteins. Hence, we observed the feature-based contribution to model performance using permutation feature importance (Fig 5). The top 10 features governing the ML performance included three Dipeptide composition (DPC) descriptors and three compositions, transition, and distribution descriptors.

**Figure 4.**
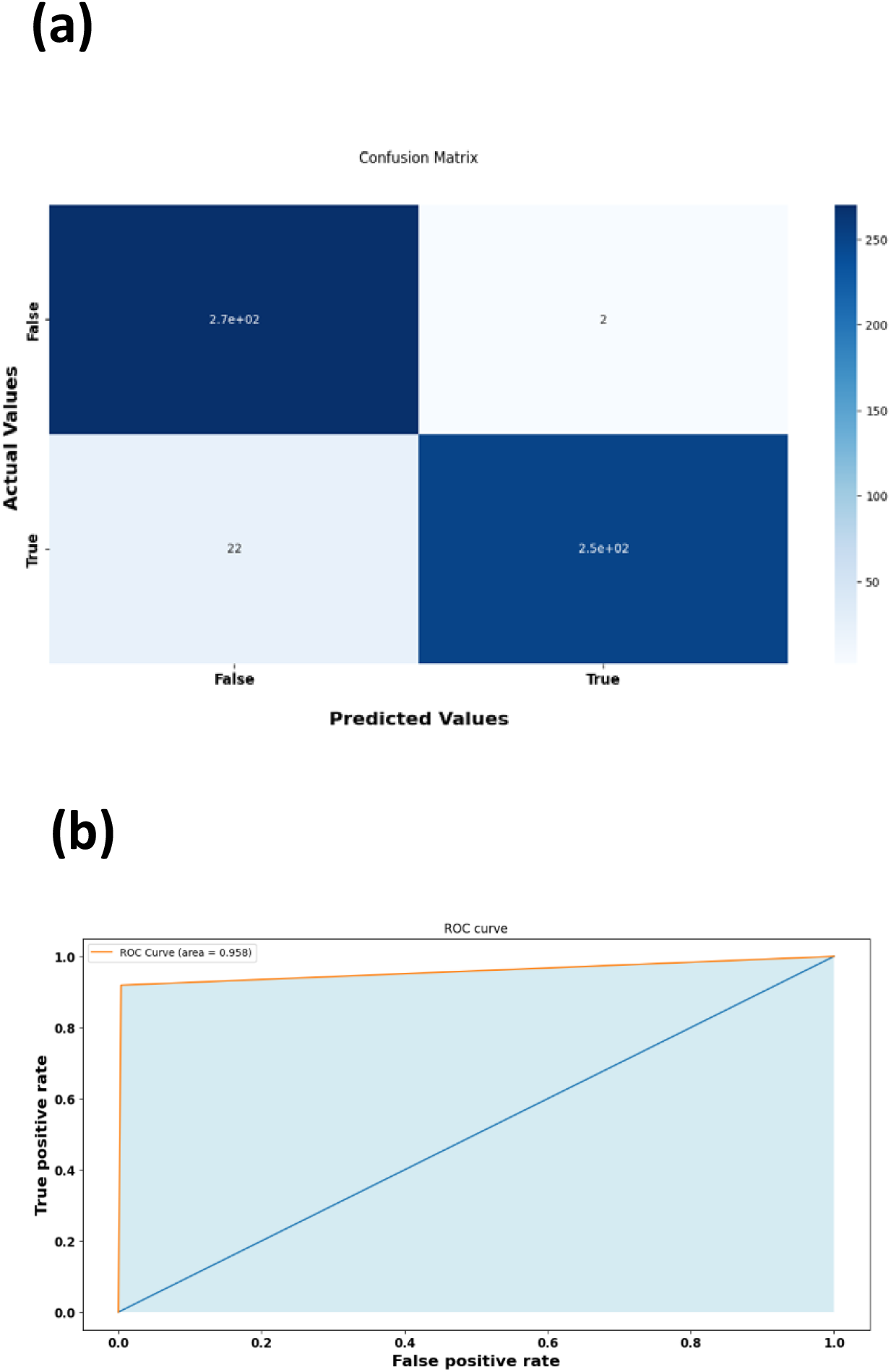
Representation of (a) accuracy, (b) Roc curve of the RF model trained with top important features

**Figure 5.**
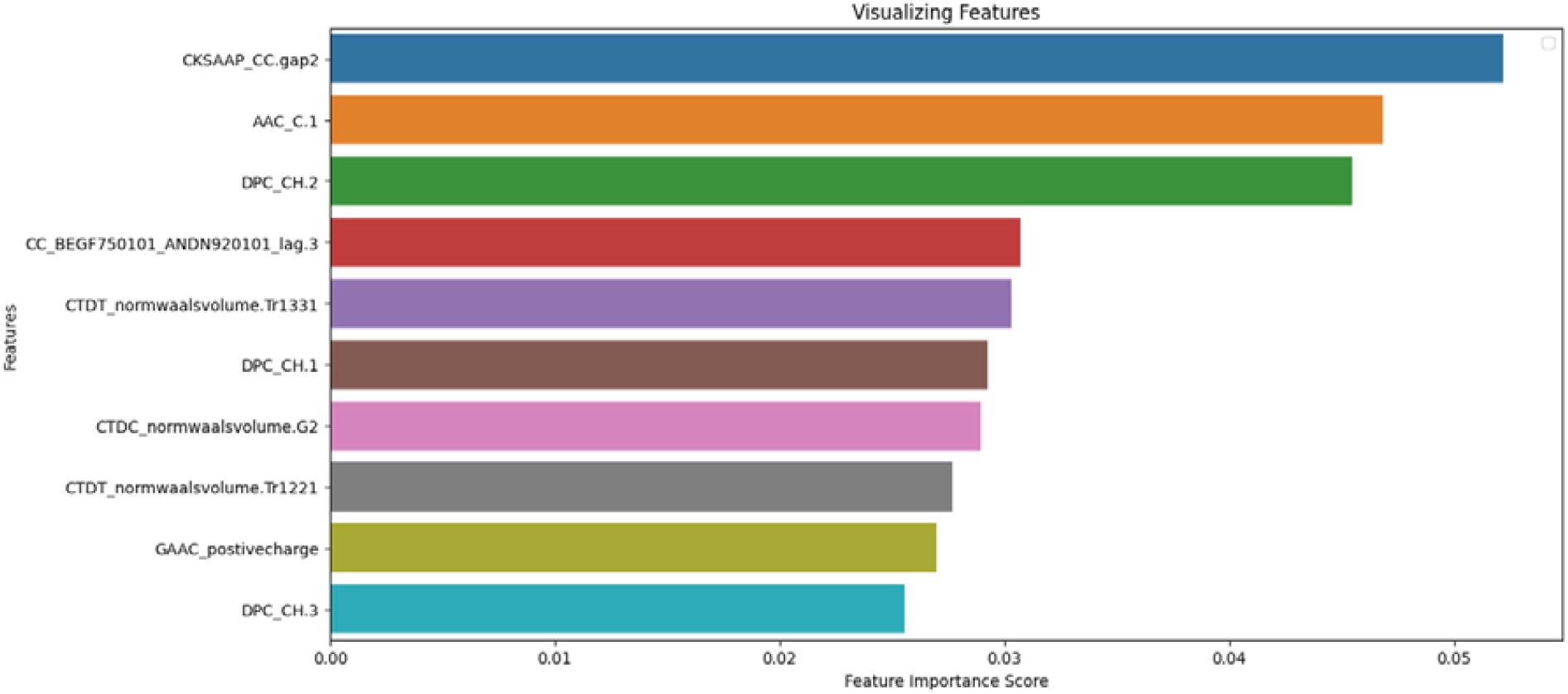
Representation of the top ten important features aiding in the decision-making by Random Forest model

## Conclusions

Nontoxic, biodegradable microbial nanowires are gaining extensive commercial attention as substitutes for synthetic NW and carbon nanotubes. With this interest and growing collection of microbial NW from diverse microorganisms, a rapid screening tool for the validation of microbial nanowires aids in reducing time-consuming experimental validation. This study created a collection of nanowire proteins through meticulous curation from available databases and literature reviews. The availability of experimentally verified nanowire protein collections incorporated into a dataset establishes a baseline for further analysis and development of a database. Using GO analysis, nanowires were found to be membranal proteins with metal ion binding domains and play a crucial role in electron transfer activity. Using ML algorithms, a model was able to predict nanowire proteins with an accuracy of 95.77%. Therefore, in this work, a model with high accuracy for the prediction of NW and its functionality was developed.

## Microbial nanowire prediction tool

The prediction tool is accessible through the web server at https://nanowire.bicbioeng.org/

## Author Contributions

DR: Design, data collection, analysis, manuscript preparation; VP: design, analysis, data curation, editing; AB: design, analysis; TDD: Webserver development. JRK: conceptualization; DRS: conceptualization; VG: conceptualization; EZG: conceptualization, guidance, design, reviewing, editing; SSD: conceptualization, design, data curation, reviewing, editing, funding.

## Figure Legends

Figure 1. Feature contributing highly towards the prediction of nanowire proteins

Figure 2. The Confusion matrix describing the actual and predicted result using Machine Learning

Figure 4. Accuracy and Roc curve of RF model trained with top 100 important features.

Figure 5. Top 10 important features aiding in the Random Forest model’s decision-making

## Supplementary Information

**Supplementary Table S1.** Accuracy and precision of different protein descriptors used for training the Random Forest machine learning model

| Protein descriptors | Accuracy | precision | recall | f1score | roc_auc |
| --- | --- | --- | --- | --- | --- |
| QSO | 0.991497 | 0.991505 | 0.991497 | 0.991498 | 0.991582 |
| GAAC | 0.954082 | 0.954083 | 0.954082 | 0.954057 | 0.953122 |
| CC | 0.989796 | 0.989901 | 0.989796 | 0.989802 | 0.990373 |
| Moreau | 0.964286 | 0.96507 | 0.964286 | 0.964339 | 0.965798 |
| AAC | 0.988095 | 0.988159 | 0.988095 | 0.988101 | 0.988486 |
| CKSAAP | 0.998299 | 0.998305 | 0.998299 | 0.998299 | 0.998113 |
| CTDC | 0.984694 | 0.984694 | 0.984694 | 0.984691 | 0.984374 |
| CTDD | 0.979592 | 0.979753 | 0.979592 | 0.979569 | 0.978375 |
| CTDT | 0.988095 | 0.988097 | 0.988095 | 0.988093 | 0.987809 |
| DPC1 | 0.998299 | 0.998305 | 0.998299 | 0.998299 | 0.998113 |
| DPC2 | 0.998299 | 0.998305 | 0.998299 | 0.998299 | 0.998113 |
| KNN | 0.988095 | 0.988159 | 0.988095 | 0.988101 | 0.988486 |
| Geary | 0.952381 | 0.952381 | 0.952381 | 0.952381 | 0.951913 |
| Moran | 0.942177 | 0.942662 | 0.942177 | 0.942244 | 0.942964 |
| SOCNumber | 0.94898 | 0.949035 | 0.94898 | 0.948996 | 0.948817 |
| AC | 0.947279 | 0.947379 | 0.947279 | 0.947304 | 0.947269 |
| AAIndex | 0.998299 | 0.998305 | 0.998299 | 0.998299 | 0.998113 |
| PAAC | 0.993197 | 0.993216 | 0.993197 | 0.993195 | 0.992792 |
| ZsCale | 0.998299 | 0.998305 | 0.998299 | 0.998299 | 0.998113 |
| TPC1 | 0.993197 | 0.993225 | 0.993197 | 0.9932 | 0.993469 |
| TPC2 | 0.994898 | 0.994905 | 0.994898 | 0.994899 | 0.995017 |
| Ctriad | 1 | 1 | 1 | 1 | 1 |

**Supplementary Figure S1.**
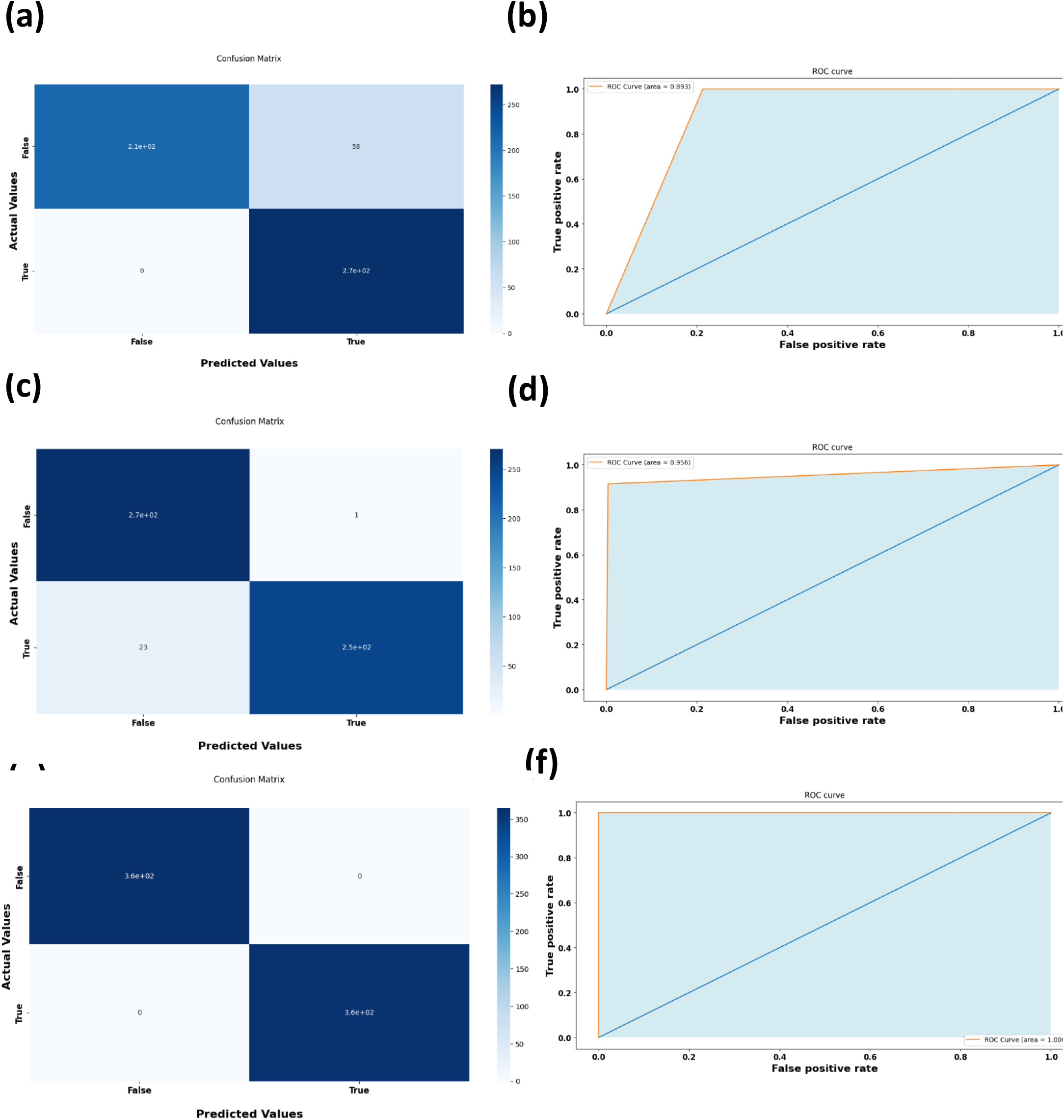
Implementation of Support vector machine (SVM), Random Forest (RF), and XGBoost for the prediction of nanowire proteins showing confusion matrix of prediction for (a) (a) SVM, (c) RF, and (f) XGBoost along with their ROC curves, respectively.

